# Woman the Hunter? Female foragers sometimes hunt, yet gendered divisions of labor are real

**DOI:** 10.1101/2024.02.23.581721

**Authors:** Vivek Venkataraman, Jordie Hoffman, Raymond B. Hames, Duncan N.E. Stibbard-Hawkes, Karen Kramer, Robert Kelly, Kyle Farquharson, Edward H. Hagen, Barry S. Hewlett, Helen Elizabeth Davis, Luke Glowacki, Haneul Jang, Kristen Syme, Katie Starkweather, Sheina Lew-Levy

## Abstract

Gendered divisions of labor are a feature of every known contemporary hunter-gatherer (forager) society. While gender roles are certainly flexible, and prominent and well-studied cases of female hunting do exist, it is more often men who hunt. A new study (Anderson et al., 2023) surveyed ethnographically known foragers and found that women hunt in 79% of foraging societies, with big-game hunting occurring in 33%. Based on this single type of labor, which is one among dozens performed in foraging societies, the authors question the existence of gendered division of labor altogether. As a diverse group of hunter-gatherer experts, we find that claims that foraging societies lack or have weak gendered divisions of labor are contradicted by empirical evidence. We conducted an in-depth examination of Anderson et al. (2023) data and methods, finding evidence of sample selection bias and numerous coding errors undermining the paper’s conclusions. Anderson et al. (2023) have started a useful dialogue to ameliorate the popular misconception that women never hunt. However, their analysis does not contradict the wide body of empirical evidence for gendered divisions of labor in foraging societies. Furthermore, a myopic focus on hunting diminishes the value of contributions that take different forms and downplays the trade-offs foragers of both sexes routinely face. We caution against ethnographic revisionism that projects Westernized conceptions of labor and its value onto foraging societies.

## Introduction

One of the most consistent and well-established empirical regularities in the study of contemporary hunter-gatherers (foragers) is the existence of a flexible sexual/gendered division of labor, within which men and women make different but complementary subsistence contributions. Men tend to spend much of their time hunting medium-to-large sized game, but often return empty-handed, whereas women spend much of their time caring for young children, gathering plant foods, and sometimes hunting small game, typically more reliable food sources [1–6]. Despite these complementary divisions of labor, which vary as a function of ecological factors [3], gender roles are not rigidly deterministic and vary with cultural context [7,8]. For instance, women make capable hunters [8,9], with reports of female participation in hunting going back centuries [10], and men regularly gather and are involved in childcare [3,11–14]. Among most foraging populations, however, women hunt rarely, and sometimes not at all. One estimate placed the frequency of women’s hunting across foraging societies at roughly 7% [15].

A recent study, published in this journal, questioned any such gendered^1^ division of labor in ethnographically known foragers. We follow Anderson et al. (2023) in referring to sex as biologically denoted traits whereas gender refers to culturally denoted traits reflecting the intersection of social norms and personal expression. Based on a survey of ethnographies that contained explicit descriptions of hunting, Anderson et al. (2023) [16] reported that women’s hunting occurs in 79% of foraging societies and that this involves large^2^ game in 33% of foraging societies. With these findings, they highlight the “…significant role females have in hunting, thus dramatically shifting stereotypes of labor…”, and go on to question the existence of any gendered divisions of labor. Other forms of labor common in foraging societies (e.g. gathering, food processing, childcare) were not analyzed by gender to bolster this claim. These claims have been widely reported in the press, e.g. [19–25], and are cited in several scientific articles [26–34].

Scientific paradigms must always be challenged. We agree that, historically, hunting and men’s labor have been over-emphasized in research among forager populations. For instance, see ref [44] for discussion of gender bias in historical datasets such as the *Ethnographic Atlas*. We applaud Anderson et al. (2023) for their willingness to challenge an orthodoxy that is often misused to justify misogyny and limit the opportunities of women [35,36]. Nevertheless, their claims about contemporary foraging societies are contradicted by the large existing literature on female hunting and the gendered division of labor. Misrepresentation of the ethnographic record devalues the many essential contributions of forager women (and men) beyond hunting, and dispenses with a century of hard-won empirical research. In this piece, we critique the claims of Anderson et al. (2023), showing that multiple methodological failures all bias their results in the same direction.

## Methodological critique of Anderson et al. (2023)

Anderson et al. (2023) coded their data at the society level. They provide no evidence that they conducted a paragraph-level analysis, which has been standard in cross-cultural research for some years [37]. It was not possible to ascertain where the relevant text was located that was used to determine women’s hunting. As a result, their analysis is not replicable. Further, their coding scheme does not account for the frequency of women’s hunting in a society in terms of effort allocated or the amount of prey acquired. Women’s hunting was coded as a binary variable and recorded as ‘present’ whether the case was a single report or habitual involvement. There was no minimum threshold for inclusion and, e.g., societies where hunting constituted only 1.2% of women’s returns [38] were coded as having women’s hunting. Beyond brief commentary on ‘opportunism’ in hunting, Anderson et al.’s (2023) results give little indication about the importance or broader societal context of women’s hunting activities.

### Sampling methodology and selection bias

Anderson et al. (2023) found evidence for women’s hunting in 50/63 (79%) of forager societies surveyed. For this estimate to be representative of actual contemporary forager diversity, the sampling procedure would need to be unbiased. Anderson et al. (2023) describe their sampling procedure as follows: they chose 391 forager societies from the D-PLACE (https://d-place.org/) ethnographic database [39] and selected that subset of 63 in which ‘explicit’ information about hunting was available. To account for autocorrelation (Galton’s problem), societies were chosen in geographically diverse locations. Little further information about sampling was provided in the text of the article. However, the senior author told the newspaper *El Pais* — for an article entitled ‘*Women have always hunted as much as men*’ — that societies were only included where sources were explicitly “detailing hunting behavior and strategies”. Studies lacking tables, statistics or details, were excluded from the sample [22]. The methods did not incorporate other statistical procedures that are common in quantitative ethnographic analysis, and the inclusion criteria were neither well described nor clearly operationalized. Given that Anderson et al.’s (2023) resulting estimate of 79% is substantially higher than previous quantitative assessments of 7% [15], it is worthwhile to consider selection bias.

“Reproduction” is defined as obtaining the same results as a given study using the same data and same methods, and “replication” as obtaining similar results using new data and similar methods as one or more previous studies [40]. We did both. We first tried to reproduce Anderson et al. (2023) by following their methods to obtain the same texts from the same database. When this failed, we tracked down most of their sources and recoded them. In the interest of reproducing the study of Anderson et al (2023) as closely as possible, we do not limit our attention to foragers. While Anderson et al. (2023) refer to all societies in their study as foragers, they did not distinguish between subsistence types in their study, labeling several horticulturalist and agriculturalist groups as foragers.

### Improper exclusion from D-PLACE

First, we explored whether Anderson et al. (2023) followed their own reported exclusion criteria. In this section of our critique, we worked with population names in the datasets as they were presented, not adjusting for potential cases of pseudoreplication. We considered, first, whether societies were included in the analysis that were not found in D-PLACE. We identified a significant number of societies in the Anderson et al. (2023) sample that were not in the D-PLACE database (which comprises the *Ethnographic Atlas*, the Binford dataset, the Standard Cross-Cultural Sample (SCCS), and the Western North American Indians dataset). Specifically, of those 63 societies with “explicit data on hunting” (p.3), 22/63 (35%) were not present in D-PLACE. Of the 50/63 societies with putative evidence of women’s hunting, 18 (36%) were not found in D-PLACE. This raises the question of how societies outside of D-PLACE were chosen, as no details were given. We found that of those 22 societies gathered from outside of D-PLACE, 18/22 (81%) had evidence of women’s hunting, according to the authors. In the absence of methodological details about how bias was avoided, this result implies biased selection.

We next considered whether there were societies among the initial 391 D-PLACE samples for which explicit descriptions of hunting *were* available, but were excluded nonetheless. Here, we found evidence that Anderson et al. (2023) did not follow their own reported selection criteria and excluded some groups that should have been included, again inflating the frequency of women’s hunting. Our attempts to acquire from D-PLACE [39] the same list of 391 societies that Anderson et al. (2023) began with did not meet with success due to a lack of sufficient information in their paper, and so we could not reproduce their analysis. Instead, to assess the possibility that some relevant ethnographies were wrongly excluded, we constrained our search to two recent large-scale and comprehensive quantitative syntheses of human foraging behavior [41,42]. We explored the overlap between these two datasets, D-PLACE, and the Anderson et al. (2023) dataset. We found 18 societies that were described in D-PLACE, in which *comprehensive explicit data* on hunting existed, *but which were excluded* from the Anderson et al. (2023) analysis. This provides evidence that the study did not follow its own exclusion criteria. Moreover, of these 18 societies (Anbarra, Arnhem, Bari, Bororo, G/wi, Gadio Enga, Kanela, Mamainde, Maya, Mekranoti, Nukak, Onge, Piro, Shipibo, Xavante, Yanomamö, Ye’kwana, Yukpa), 9 (G/wi, Nukak, Onge, Yanomamö, Arnhem, Shipibo, Mamainde, Bari, and Maya) reported explicitly that women did not hunt, and in the other 9 only men’s hunting was mentioned (see footnote 5). This again raises the possibility of biased selection and again serves to artificially inflate estimates of women’s hunting. A comprehensive re-analysis of the entire D-PLACE database would likely reveal more cases of improper exclusion.

### Selective inclusion of non-D-PLACE sources

We next considered Anderson et al.’s (2023) inclusion of non-D-PLACE sources. Koster et al. [41] selected datasets that characterized hunting across the lifespan, and Kraft et al. [42] selected studies that characterized subsistence contributions in-depth at the societal level, partly to characterize gender-based differences in foraging. As these studies summarize some of the most authoritative datasets on forager subsistence behavior, they constitute perhaps the best explicit descriptions currently available. Between these two publications, there were 60 unique societies. Of these, Anderson et al. (2023) included 18 in their study (!Kung, Ache, Agta, Aka, Baka, Batek, Bofi, Cree, Efe, Gunwinngu, Hadza, Hiwi, Inuit, Martu, Mbuti, Pume, Punan, Tsimane), and, of these, 15 were coded as positive for the occurrence of women’s hunting. 14 of these societies were contained in D-PLACE. This leaves 42 societies with comprehensive descriptions of subsistence behavior, including hunting, that they did not investigate.

Of the 60 unique societies between Koster et al. [41] and Kraft et al. [42], 34 were contained in D-PLACE, and 26 of these societies were not (Achuara, Batek, Bofi, Dolgan, Etolo, Hiwi, Inujjuamiut, Kaul, Lufa, Machiguenga, Maku, Maring, Mvae, Mayangna, Nen, Nimboran, Nuaulu, Nunoa, Quichua, Tatuyo, Tsembega, Tsimane, Waorani, Wola, Wayana, Yassa). Given that Anderson et al. (2023) themselves appear to have sought sources outside of D-PLACE, it is not clear why many of the most comprehensively described societies were not chosen. In these populations, based on published work [41–43], women’s hunting occurs rarely or is not reported at all. For example, Koster et al (2020)[41] presented data on 40 small-scale societies. The data are largely unbiased toward the description of female hunting (J. Koster, *pers comm*). Only 3/40 societies (Baka, Tsimane, Martu) showed evidence for women’s hunting. Similarly, the SI of Kraft et al. (2021)[42] has detailed information on the source data, including gendered aspects of foraging.

Of course, the absence of evidence does not constitute evidence of absence, and it is always possible that women’s hunting was underreported. However, given that the Koster et al. [41] and Kraft et al. [42] datasets drew on studies where long periods of fieldwork were conducted, in the first case with particular emphasis on hunting, it is specifically these societies where evidence of women’s hunting is least likely to have been missed. This shows that even if Anderson et al. (2023) had strictly followed their own exclusion criteria, by limiting their search to D-PLACE, they would have missed a great deal of highly pertinent evidence that runs contrary to their conclusions.

We acknowledge that this is not a precise reproduction, as Anderson et al. (2023) did not provide enough detail in their methods to make this possible. Given the challenges of working with large ethnographic databases, we sympathize with the efforts of Anderson et al. (2023) and do not believe they were intentionally cherry-picking. Our point is that selecting different societies would have resulted in a far different (lower) estimate. Moreover, adding more societies to the denominator of Anderson et al.’s (2023) frequency calculation would change their estimate drastically. This is without examining the more than *300 other societies* that were in their original (unsourced) sample, in some of which hunting is likely described in detail. Even putting aside their numerous coding errors (see next section), Anderson et al.’s (2023) estimate of 79% is unreliable and probably heavily inflated. A full analysis of several hundred ethnographic reports would be necessary to arrive at a defensible estimate.

### Coding errors

Second, we thoroughly re-examined the source ethnographies for the 63 societies Anderson et al. (2023) claimed as providing explicit evidence of women’s hunting. Their data are included in their Table 1 and in the associated Supporting Information data file. We describe several issues with this dataset, the coding decisions, and the inferences that were subsequently drawn.

**Table 1:** Our recoding of women’s frequencies of hunting small-medium and large game. Societies in bold were coded by Anderson et al. (2023) as women hunting small or medium game (left) or large game (right). Bold entries in the *No evidence, Never*, and *Rarely* rows are thus inconsistent with our recoding. The selection of societies in Anderson et al. (2023) is heavily biased, so our recoded values are not representative of female hunting in foraging societies.

| Frequency | Women hunt small-medium game (<45kg) | Women hunt large game (≥45 kg) |
| --- | --- | --- |
| No evidence | [23]: **Ganij, Kalaallit, Maidu, Northern Ache, Tiwi**, Central Eskimo, Gunwinygu [sic], Iroquois, Karajarri, Kaurareg, Kikuyu, Larrakia, Mescalero Apache, Missinippi Cree, Nootka, Northern Ojibwa, Punan, Rainy River Ojibwe, Savanna Pumé, Tamang, Tolowa, Tongva, Yamana | [32]: **Matses, Mbuti, Nootka**, Adnjamatana, Australian Mardudjara, Batak, Batek De', Bofi, Ganij, Gunwinygu [sic], Hadza, Iroquois, Karajarri, Kaurareg, Kikuyu, Larrakia, Maidu, Maniq, Northern Ache, Punan, Rainy River Ojibwe, Savanna Pumé, Sua (Tswa), Tamang, Tolowa, Tongva, Torres Strait Islanders, Tsimane, Walbiri, Wopkaimin, Worrorra, Yamana |
| Never | [2]: **Torres Strait Islanders**, Jahai | [6]: **Efe, Tasmania**, Baka, Basin-Plateau, Fish Lake Valley North Paiute, Jahai |
| Rarely | [6]: **Bofi, Hadza, Lardil**, Hiwi, Iñupiaq, Kaiadilt | [16]: **!Kung San, Australian Martu, Bakola, Belcher Island Eskimo, Iñupiaq, Missinippi Cree, Northern Ojibwa**, Aka, Alyawara, Cree, Hiwi, Inuit, Ju/hoansi, Kaiadilt, Lardil, Tiwi |
| Sometimes | [11]: !Kung San, **Aka, Baka, Bambote, Batek De', Ju/hoansi, Sua (Tswa), Tsimane**, Ainu, Efe, Mono Lake Northern Paiute | [6]: **Ainu, Central Eskimo, Mescalero Apache**, Bambote, Kalaallit, Mono Lake Northern Paiute |
| Frequently | [21]: **Adnjamatana, Alyawara, Australian Mardudjara, Bakola, Basin-Plateau, Batak, Cree, Fish Lake Valley North Paiute, Gosiute, Inuit, Tasmania, Walbiri, Wopkaimin, Worrorra**, Agta, Australian Martu, Ayta, Belcher Island Eskimo, Maniq, Matses, Mbuti | [3]: **Agta, Ayta**, Gosiute |

Coding ethnographic texts on a particular topic, such as women’s hunting, is difficult because the relevant material is typically short (sometimes one sentence or less) and often ambiguous.

Coding decisions almost always require judgment calls that may differ among coders. It is therefore essential that studies of the ethnographic record identify the specific texts used for each coding decision. Because Anderson et al. (2023) did not conduct a paragraph-level analysis, we do not know what evidence was used from source ethnographies to support their coding decisions. Accordingly, we re-coded the source ethnographies used by Anderson et al. (2023). We selected text fragments that were most relevant for coding the presence of women’s hunting, the frequency of hunting, and the size of prey hunted. In our re-analysis, for a case to count as positive evidence for women’s hunting, it must involve ‘active’ participation by women (not spiritual or ritual purposes).

Although other sources contained relevant information about women’s hunting in these societies that was not consulted by Anderson et al. (2023) (see below), we confined our re-analysis to *only* the ethnographic sources - and sources used therein - used by Anderson et al. (2023). Thus, our goal was not to conduct an authoritative analysis of women’s hunting, nor to conduct a paragraph-level analysis of Anderson et al.’s (2023) 63 societies. Rather, our goal was to investigate how Anderson et al. (2023) applied their coding scheme to their sources. We found that they applied it unevenly and insufficiently, and in many ethnographies they cite there are explicit statements that contradict their coding. Our re-coding of the 63 societies contained in Anderson et al. (2023) can be found in Table S2, and serves as the basis for the following points.

### Subsistence and population selection

The first issue pertained to subsistence and population selection. Though nominally focused on hunter-gatherer (forager) subsistence, Anderson et al. (2023) included some horticultural and large-scale agricultural societies in their dataset, incorrectly described as foragers. Mixed-subsistence horticulturalists (H, HH) and agriculturalists (A) mis-coded as foragers in their list of 63 societies include the Kikuyu (A), Wopkaimin (HH), Iroquois (A), Tsimane (H), and Matses (HH). This is likely because D-PLACE does not classify societies by subsistence type, but rather by the relative contribution of food sources to the diet. Though all anthropologists acknowledge that these historical subsistence labels are imperfect and may not reflect current subsistence behavior, it is typical in cross-cultural research to define hunter-gatherers as peoples who depend on foraging for 90% of their diet [6]. Such labels also acknowledge populations’ current land rights or lack thereof [45].

### References to secondary literature

Second, we found that many references were taken from the secondary literature. Drawing on secondary sources is problematic because they are often divorced from ethnographic context and strip relevant information that might have been present in the source ethnography. Specifically, 15/63 cases were secondary references, and 14/63 cases referenced a single nearly 30-year-old secondary source [7]. Significant discrepancies are apparent between Table 1 of Anderson et al. (2023) and their Supporting Information spreadsheet, with 31/63 cases having different references. Also, 15/63 cases on the Supporting Information table did not have a reference listed at all. Also in 15/63 cases, Anderson et al. (2023) cited a source with evidence that comes from at least one other source that they do not reference. We refer to these as undisclosed secondary sources (USS; see Table 1).

### Inconsistent coding

Third, coding was inconsistent. For example, the Jahai of Malaysia were coded as not having female hunters based on an explicit statement in van der Sluys [46] that ‘women never hunt.’ Yet this is the same type of statement found for the !Kung, which Anderson et al. (2023) coded as a society in which women do hunt: “… women are totally excluded from hunting. Women never participate in a !Kung hunt…” [47]. Similarly, Lee (1979: 235)[48] wrote “Women do not hunt.”

### Insufficient search for source material

Fourth, though Anderson et al. (2023) investigated each society “by searching through the original references cited in D-PLACE [39], Binford [49], and by searching digitized databases and archives,” there are instances in which well-known authoritative sources were not consulted. For example, Anderson et al. (2023) coded the Batek of Malaysia as having female hunters based on Endicott [50]. However, a more recent book by the same author provides quantitative information on female contributions. Endicott & Endicott [51] wrote: “Still, women procured 2 percent by weight of the animals hunted by nonblowpipe methods and 22 percent of all bamboo rats.” Women procured no animals using the blowpipe (Table 4.1, p. 76)[51]. The !Kung were also coded by Anderson et al (2023) as having female hunters. Yet in her famous ethnography *Nisa: The Life and Words of !Kung Woman*, Shostak (1982:220)[52] wrote: “!Kung women cannot be considered hunters in any serious way…” A similar case prevails for the Tsimane horticulturalists of Bolivia. The authors cite Medinaceli and Quinlan [53], but they ignore a recent case study on Tsimane women hunting [54].

### Pseudoreplication

The fifth issue concerns pseudoreplication, in which the same case is counted more than once. This leads to inflated and inaccurate estimates. There are several examples. The !Kung and Ju/hoansi are treated as independent points, but these terms refer to the same population [48] . The same holds for the Agta and Ayta of the Philippines [9]. Moreover, the Efe, Sua, Mbuti (BaMbuti), and Bambote, and the Mardujara and Martu (Martu), are each counted independently despite being members of closely related groups [55,56]. We recognize that these errors by Anderson et al. (2023) are not deliberate. Indeed, in at least one case it may be valid to count these as independent data points. The Efe and Mbuti live nearby but are known to have divergent hunting strategies. The Efe are traditionally bow hunters, whereas the Mbuti are traditionally primarily net hunters [57]. However, due to the potential for cultural autocorrelation to inflate the frequency of women’s hunting, such decisions should be acknowledged and justified.

### No mention of women hunting in source material

The sixth issue is that, for some references, we could not track down any mention of women’s hunting at all in the source text (e.g., Ganij). In another case, hunting was mentioned, but it did not involve women (Lardil). A further case described trapping snares being made of women’s hair, but women are not otherwise described as being involved in hunting labor (Maidu). In several instances (the Lardil, Torres Strait Islanders, Inuit, and Kalaallit) fishing was coded as hunting, a distinction that was not made clear in the manuscript text even though these categories are customarily distinguished by anthropologists [6].

### Hunting is not a binary phenomenon

The seventh issue is that Anderson et al. (2023) conflate *any* mention of women’s hunting with evidence of women being consistently involved in hunting, especially of large game. For example, Ruth Landes’ [58] study of Ojibwa women is cited as evidence of women hunting. Indeed, Landes describes a few cases of women who chose to be hunters. However, she also states “those women who cross the occupational line and take up men’s work, do so casually, under the pressure of circumstances or of personal inclination….[Other] Women regard them as extraordinary or queer.” That is, Landes is noting that these women are very much the exception rather than the rule (Landes 1938: 136). Likewise, for the Central Eskimo (Inuit), Boas describes two instances of women hunting, one a secondary report from which he obtained from John Rae’s 1845-46 account of his travels. Rae’s account, which Boas recounts in its entirety, is a description of women hunting sun-bathing seals opportunistically with clubs (since they don’t carry spears). The other is an account of women assisting in a communal hunting effort of breathing-hole sealing, in which the women (and children) guard breathing holes and scare the seals away, forcing them to surface at a hole where a man is stationed with his spear ready. Again, though, Anderson et al. (2023) do not relate Boas’s comments that speak to the general pattern of behavior: For a typical day, the men go hunting, “meanwhile the women, who stay at home, are engaged in their domestic occupations, mending boots and making new clothing …” (Boas, 1889)(1964: 154)[59]. And, “the principal part of the man’s work is to provide for his family by hunting, i.e., for his wife and children….The woman has to do the household work, the sewing, and the cooking. She must look after the lamps, make and mend the tent and boat covers, prepare the skins, and bring up young dogs.” (Boas, 1889)(1964: 171-2)[59].

#### A re-analysis of Anderson et al.’s (2023) ethnographic sources

In light of these issues, we re-coded three of Anderson et al.’s (2023) variables using the same ethnographic sources they cite (Table 1 and Fig 1). In place of Anderson et al.’s (2023) binary hunting variable, we coded the frequency of women’s hunting in each society based on contextual information (Table 1). Our five hunting frequency categories were: no evidence for women hunting, evidence that women did not hunt, and evidence for women hunting rarely, sometimes, or frequently. We did this separately for two prey sizes. Anderson et al. (2023) coded four prey size categories: small game (1), medium game (2), large game (3), and all (4). The criteria for distinguishing small, medium, and large game were not specified or operationalized, and no standard body-size cut-off was used. In some cases, prey size was determined indirectly. For example, in some instances, prey species were coded as ‘large game’ based on being struck with large sticks and machetes. As large sticks and machetes can be used to kill prey of a range of body sizes, we find this criterion problematic. We followed a standard cut-off used in zooarchaeology and paleontology, adopting the criteria of small-medium game (<45 kg) versus large game (≥45 kg) [17,18]. Accordingly, we had two women’s hunting variables: the frequency of hunting small-medium game, and the frequency of hunting large game. We also re-coded the primary mode of subsistence for each society. See the Appendix for our detailed coding rubric and the inter-rater reliability of our independent coders.

**Fig 1:**
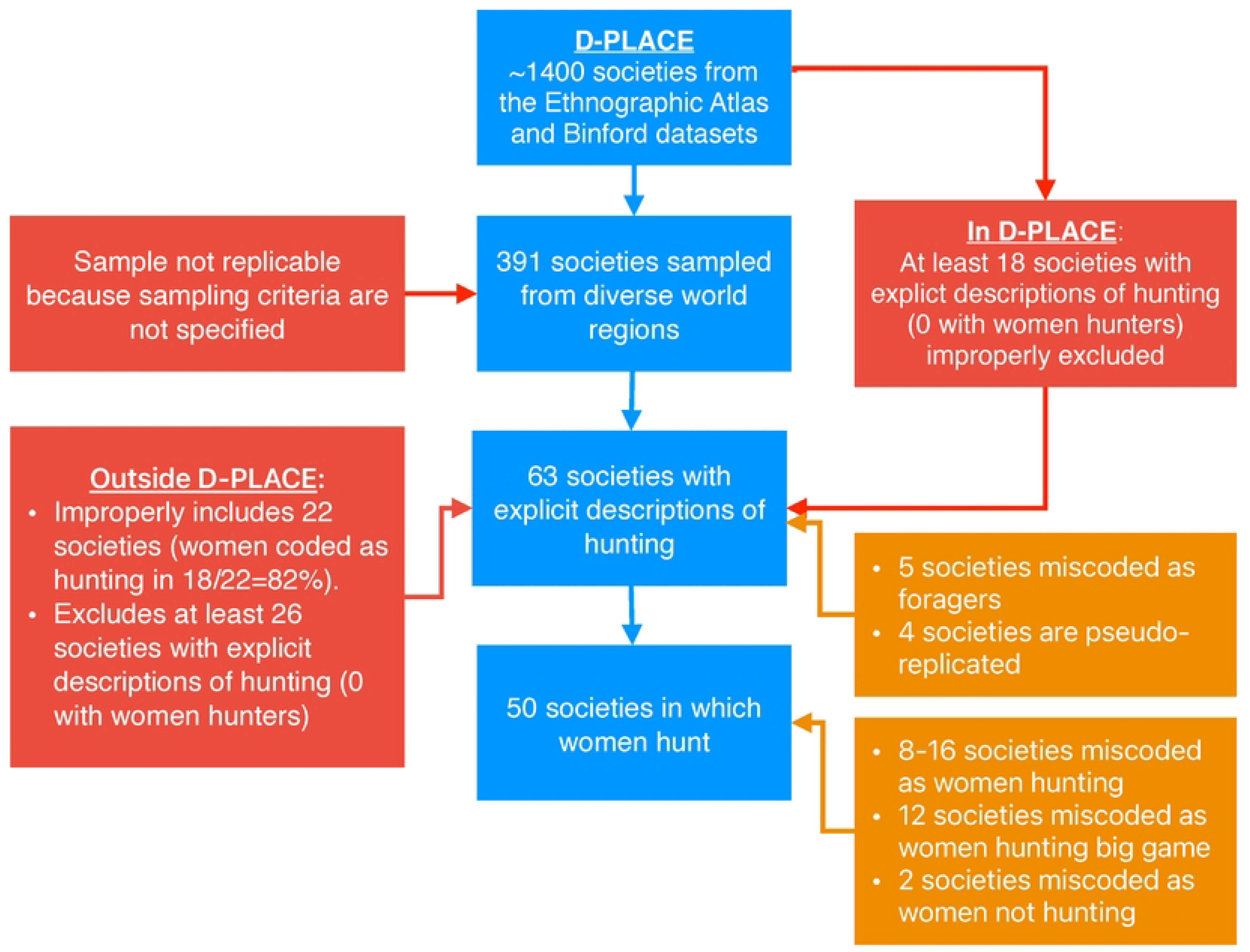
Diagram highlighting issues with the sampling methodology and coding of Anderson et al. (2023). Blue boxes indicate data sampling and coding by Anderson et al. (2023). Red boxes describe sampling errors. Orange boxes describe coding errors. Nearly two dozen societies were drawn from outside D-PLACE without explanation, and these were heavily biased toward positive reports of female hunting. At least 18 societies were present in D-PLACE that had explicit descriptions of hunting but were not included in the analysis of Anderson et al. (2023); none of these describe women’s hunting (see footnote 5). We also identified at least 26 societies outside of D-PLACE with explicit descriptions of hunting. Women’s hunting is uncommon or absent in these societies. Anderson et al. (2023) appear to have biased their inclusion of non-D-PLACE societies in their analysis toward societies in which women hunt. Taken together, improperly including societies biased toward women’s hunting and improperly excluding societies with little or no evidence of women’s hunting likely led to sampling bias. A definitive accounting of the proportion of forager societies in which women hunt must await a future study with a rigorous and unbiased sampling strategy.

We then compared the binary Anderson et al. hunting variable with our two hunting frequency variables. Of the 50 societies that Anderson et al. (2023) coded as women hunting, we found that in 8 of them, there was either no evidence for women hunting or evidence against women hunting. In a further 8, there was only evidence that women hunted rarely. Hence, from 8/50 (16%) to 16/50 (32%) of the societies that Anderson et al. (2023) coded as women hunting were false positives. We also found that 2/13 (15%) of the societies that Anderson et al. (2023) coded as no women hunting in fact did have evidence of women hunting, i.e., were false negatives. See Fig 2.

**Fig 2:**
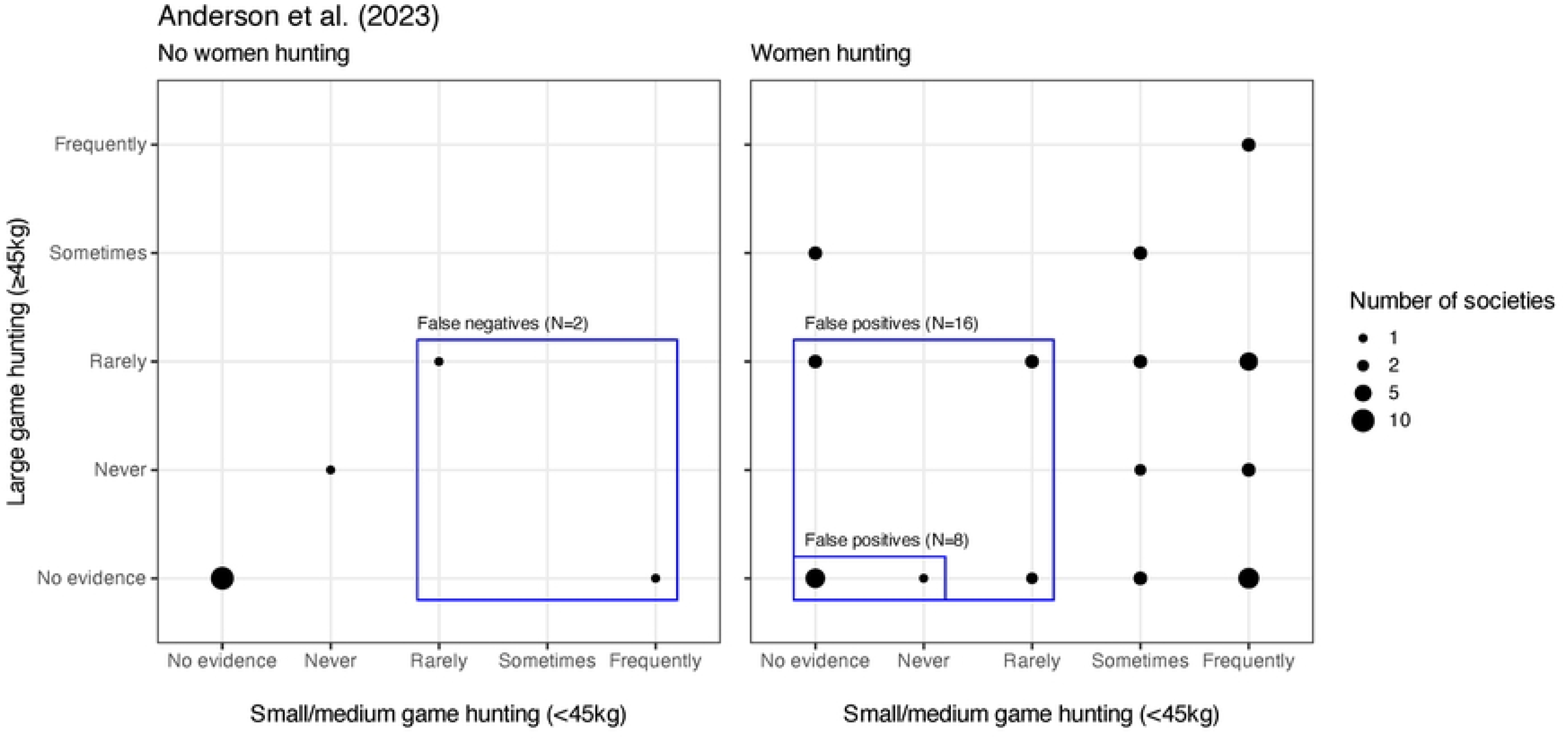
Comparison of Anderson et al.’s (2023) coding (left and right panels) to our re-coding (x and y axes). Dot size indicates the number of societies with that combination of women’s small and large game hunting frequency. **Left**: Societies that Anderson et al. coded as women do not hunt. There were 2 societies for which we found some evidence of women hunting (blue box, false negatives). **Right**: Societies that Anderson et al. coded as women hunt. There were 8 societies for which we either found no evidence that women hunted, or evidence that women did not hunt, and 16 societies for which we either found no evidence that women hunted, or evidence that women’s hunting was either rare or absent (blue boxes, false positives). See also Table 1 and Table S1. The selection of societies in Anderson et al. (2023) is heavily biased, so our recoded values are not representative of female hunting in foraging societies.

We also re-assessed the claim that a third or more of the surveyed societies showed evidence of women participating in big-game hunting by comparing the Anderson et al. (2023) prey size variable with our two hunting frequency variables. Of the 63 societies with ‘explicit evidence of hunting’, Anderson et al. (2023) located prey sizes for 45, 15 of which they coded as hunting big game (33%), and 2 as hunting game of all sizes, for a total of 17/45 (38%) societies as women being involved in large-game hunting. In our re-coding of women’s large game hunting frequency for these 17, we instead found No evidence: 3/17 (18%); Never: 2/17 (12%); Rarely: 7/17 (41%); Sometimes: 3/17 (18%); and Frequently: 2/17 (12%). Thus, less than a third of those originally coded involved regular female large-game hunting. Of the cases that do involve large-game hunting by women, in multiple instances the ethnographer explicitly stated that it was *very rare*. Some only described cases of women hunting large game alone after the death of their husbands. Other cases pertained to communal whale hunting, or involved dogs. Some occurred in the context of husband-wife pairs where women contributed to hunting success indirectly. Some cases involved firearms, a uniquely modern technology that was unavailable during human prehistory.

In our own re-coding of the frequency of big game hunting in all 63 societies (i.e., in the 17 that Anderson et al. coded as big game hunting, as well as in the 46 that they coded as no big game hunting, and ignoring pseudo-replication) we found the following, No evidence: 32/63 (51%); Never: 6/63 (10%); Rarely: 16/63 (25%); Sometimes: 6/63 (10%); and Frequently: 3/63 (5%). Thus, in only 9/63 (14%) societies in Anderson et al. (2023) could women be said to “regularly” hunt big game (but two of those 9, the Agta and Ayta, are the same population). See Table 1 and Fig 3.

**Fig 3:**
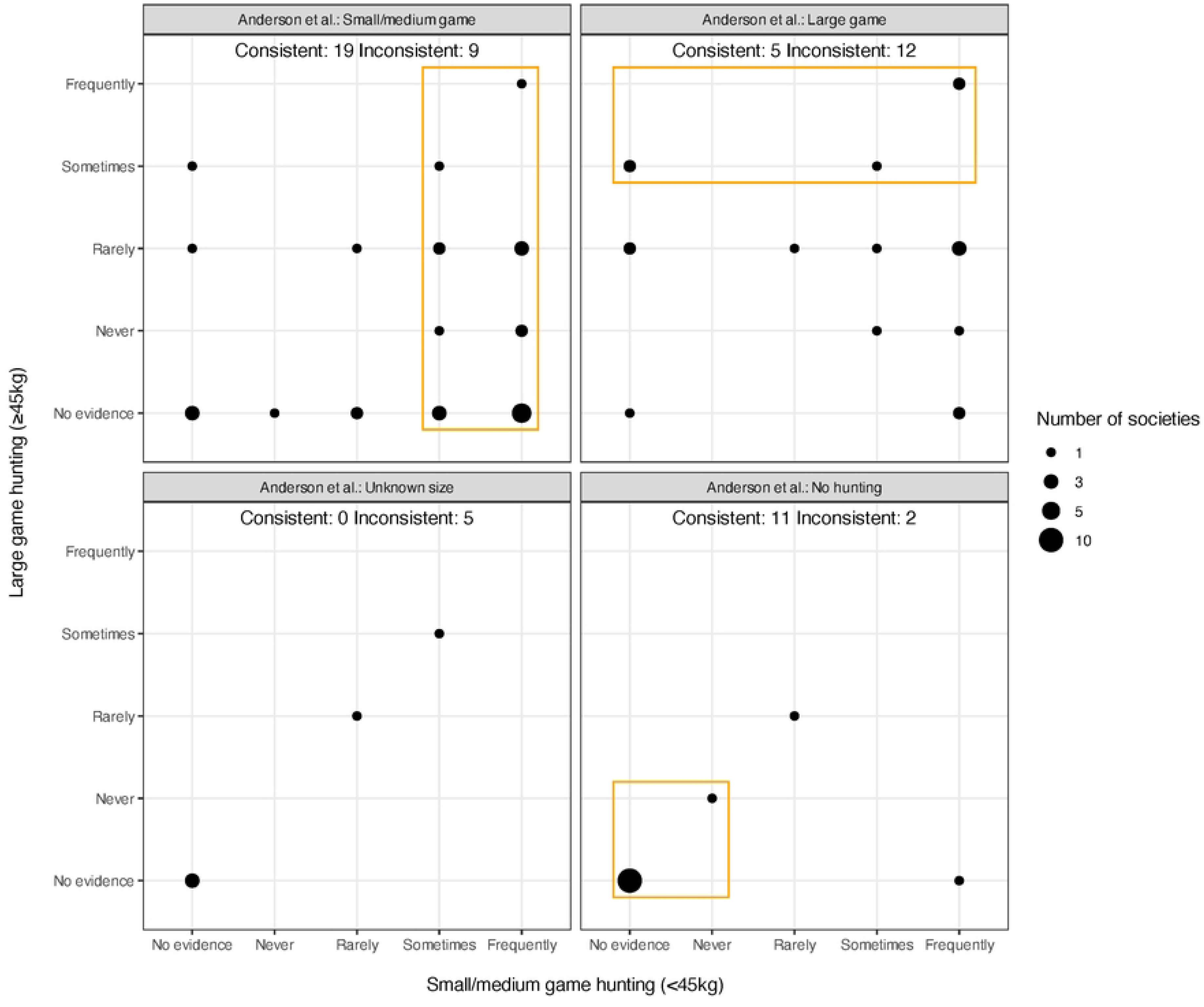
Comparison of Anderson et al. (2023) prey size coding (the four panels) with our prey size re-coding (x and y axes). Dot size indicates the number of societies with that combination of women’s small and large game hunting frequency according to our coding. For comparison with our coding, we combined the Anderson et al. small (1) and medium (2) categories. Societies inside and outside the orange boxes are those where the Anderson et al. coding of prey size is consistent and inconsistent with ours, respectively. Anderson et al. coded two societies as women hunting both small-medium and large game, which we placed in the “Large game” panel (upper right). We categorized frequent or occasional hunting of a prey size as consistent with the claim that women hunted that prey size, and inconsistent otherwise. The selection of societies in Anderson et al. (2023) is heavily biased, so our recoded values are not representative of female hunting in foraging societies.

Due to the sampling bias and insufficient search for source material in Anderson et al. (2023), our revised estimates of the frequency of women’s hunting in their dataset are *not* to be taken as representative of foraging societies. Our point, rather, is that (1) the Anderson et al. (2023) data, even though likely heavily biased toward societies in which women hunt, nevertheless only support a much lower estimate of frequent hunting by women, and (2) the estimates can vary wildly depending on inclusion criteria and coding decisions.

## Conclusion

We agree with Anderson et al. (2023) that we should dispel the categorically false notion that forager women do not hunt or are unable to hunt, and we thank them for bringing this important topic to the public forum. Though we appreciate their intent, Anderson et al. (2023) do not overturn current consensus views on gendered divisions of labor among contemporary foragers, which are based on a substantial body of empirical evidence. Like Anderson et al (2023), we acknowledge this body of evidence is significantly filtered through the lenses and experiences of 20th-century ethnographers who may have had their own biases and interests but were likely less aware of their positionality and its impact on their subsequent ethnographic descriptions.

We have outlined several conceptual and methodological concerns with Anderson et al.’s (2023) analysis. Specifically, Anderson et al.’s (2023) analysis is not reproducible because their sampling criteria are not clear and 35% of the societies in their sample do not come from D-PLACE, the database they claim was the source of all the societies in their sample. Moreover, these 35% were heavily biased toward societies that they coded as ones in which women hunt.

Many other societies with extensive information on hunting are also not in D-PLACE yet were not included in their analysis, and authoritative sources on hunting in the societies in the Anderson et al. (2023) sample were not consulted. Additionally, there are at least 18 societies in D-PLACE with information on hunting that were inexplicably omitted from their analysis, none of which provide evidence for women hunters.

Finally, there were numerous coding errors. Of the 50/63 (79%) societies that Anderson et al. (2023) coded as ones in which women hunt, for example, our re-coding found that women rarely or never hunted in 16/50 (32%); we also found 2 false negatives. Overall, we found evidence in the biased Anderson et al. (2023) data set that in 35/63 (56%) societies, women hunt “Sometimes” or “Frequently”. Moreover, compared to the 17/63 (27%) societies in which women were claimed to hunt big game regularly, our re-coding found that this was true for only 9/63 (14%). A precise estimate of women’s hunting in foraging societies must await a future thorough and unbiased analysis of the ethnographic record (see, e.g., [10]), but it is certainly far less than the Anderson et al. (2023) estimate and is very unlikely to overturn the current view that it is relatively uncommon.

The fundamental issue is that women’s hunting is not a binary phenomenon, and treating it as such, especially with a very low threshold for classifying a society as one in which women hunt, obfuscates gendered divisions of labor within groups. Anthropologists have long recognized that the nature of cooperation in foragers is complex and multi-faceted, and women’s and men’s subsistence activities play important and often complementary roles. Moreover, women’s hunting has been studied for decades, and anthropologists have a good understanding of when and why it occurs. Yet, to focus on hunting at the expense of other critical activities - from gathering and food processing, to water and firewood collection, to the manufacture of clothing, shelters, and tools, to pregnancy, childbirth, nursing, childcare, and healthcare, to education, marriages, rituals, politics, and conflict resolution - is to downplay the complexity, and thereby the importance of women’s roles in the foraging lifeway. To build a more complete picture of the lives of foragers in the present and the past, it serves no one to misrepresent reality. In correcting the misapprehension that women do not hunt, we should not replace one myth with another.

## Supporting information

### S1 Appendix

#### Re-coding of Anderson et al. (2023)

Re-analysis of the 63 societies coded by Anderson et al. (2023), i.e., those for which they claim to have found explicit descriptions of hunting. Our data file included the following five fields from the Anderson et al. (2023) data file (however, our R script downloads that data file directly from the the PLoS ONE website):

- **Society**: as found in Anderson et al. (2023).
- **Subsistence**: as stated by Anderson et al. (2023).
- **Reference**: as stated in Anderson et al. (2023), along with a comment. USS = undisclosed secondary source; this refers to the case in which Anderson et al. (2023) cited a source with evidence that comes from at least one other source that is not directly referenced by Anderson et al. (2023). Confirmation source = source(s) from Anderson et al. (2023) that we found most relevant to confirming women hunting.
- **Prey size category**: as coded by Anderson et al. (2023): small (1), medium (2), large game (3), all (4).
- **Presence of women’s hunting**: as coded by Anderson et al. (2023).

We used the following rubric to recode each row in the Anderson et al. (2023) data set. The rubric was developed as follows. First, VV, JH, and KF developed a coding rubric and coded all 63 societies. This rubric was used by rater 2 (SL) to independently recode the hunting frequencies of small and large game in all 63 societies. She found this initial rubric to be insufficiently detailed to confidently code all societies. Based on feedback from SL, the rubric was updated by VV, JH, and KF. Rater 3 (KS) then independently recoded the hunting frequencies of small and large game of all 63 societies using the following final rubric:

#### Subsistence

Re-coding of the subsistence type of Anderson et al. (2023), according to D-PLACE or other primary literature sources.

- HH = Hunter-Horticulturalists
- A = Agriculturalists
- H = Horticulturalists
- HG = Hunter Gatherers
- F = Fishers

#### Information from references

Within the source ethnographies used by Anderson et al (2023), we selected text fragments that appeared to us the most relevant for coding the presence of women’s hunting, the frequency of hunting, and the size of prey hunted. While other sources contained relevant information about women’s hunting in these societies that was not consulted by Anderson et al. (2023), we confined our re-analysis to *only* the ethnographic sources - and sources used therein - used by Anderson et al. (2023). For a case to count as positive evidence for women’s hunting in our re-coding, it must involve ‘active’ participation by women (not only spiritual or ritual purposes).

#### Species & body size

Refers to prey specifically hunted by women. When prey pursued not specifically referred to, all potential prey listed. Species names taken from source ethnography; various online sources used to find average body weights. Most paragraphs contained information about prey species pursued. When this was not the case and thus no information about body size was available, we sourced potential prey species from online or ethnographic sources. These sources are listed. In cases where women are clearly not hunting based on the evidence, prey species not listed.

#### Hunt small-medium game? (<45kg), Hunt large game?(>=45kg)

For these columns, we assessed the frequency of hunting. In cases where the body size of prey straddles the 45 kg cutoff, both columns were coded. When conflicting information is present between or within paragraphs (see the case of the !Kung, in which it is said ‘women do not hunt’, but in the same paragraph it is stated that they participate as beaters), we consider the concrete statements to overrule the general statement, thus favoring positive coding of women’s hunting.

- Frequently: Participation in hunting with some kind of regularity, including seasonal behavior. Example phrases include ‘frequently’, ‘always’, ‘common’, ‘often’, or similar.
- Sometimes: Participation in hunting occurs less often than frequently but more often than rarely. Example phrases include ‘occasionally’, ‘sometimes’, or similar. When cases do not give any indication of frequency but state ‘women hunt’ or the statement involves the word ‘do’, we coded this as ‘frequently’, as such definitive statements are considered to refer to habitual behavior. This is downgraded to ‘sometimes’ if there is other evidence suggesting that the behavior does not qualify as frequent (see the case of the Sua).
- Rarely: Participation in hunting occurred one time or rarely. Alternatively, it was based on hearsay, indicating the ethnographers themselves never observed it (suggesting rarity). This was also coded for inferred cases (e.g. authors state that net-hunting involves women, and that net-hunting involves capture of given species, but not explicitly stated that women capture given species). Phrases such as accidental, opportunistic, and chance fell under this coding.
- Never: Participation in hunting by women said explicitly to not occur, or that only men hunt.
- No evidence: No mention of women hunting, but cannot be definitively excluded. When only small or large game are mentioned, the unmentioned size class is classified as ‘no evidence.’

#### Pseudoreplication

Societies with different names, such as !Kung San and Ju/hoansi, coded as different societies in Anderson et al. (2023), but are in fact the same society.

#### Summary

Our reasons for our codings of women’s small and large game hunting frequency.

#### Inter-rater reliability

We evaluated inter-rater reliability using agreement plots and Bangdiwala’s *B* statistic [60,61] separately for small and large game, and for raters 2 and 3 vs. the original coding. Agreement plots depict the degree of agreement and disagreement for each category. Bangdiwala’s *B* [61] is the chance-corrected degree of agreement computed from the contingency table of the ratings of two raters:

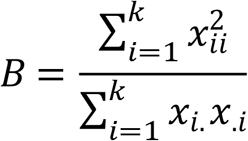

where *x*_*ii*_ are the values on the diagonal (i.e., the counts of agreement), and *x*_*i*._ and *x*_.*i*_ are the sums of row *i* and column *i*, respectively. The denominator is therefore the maximum possible agreement, given the marginal totals. Values of *B* ≥ 0.81 are interpreted as “almost perfect” agreement [60].

The weighted Bangdiwala statistic [61] for rater 2 agreement for small game was B = 0.73, and for rater3 was B = 0.8. See Fig. S1.

Fig. S1: Small game agreement plots for the original ratings (x-axis) vs. rater 2 (y-axis, left) and rater 3 (y-axis, right). The larger outer white rectangles represent the maximum possible agreement for each category, the inner black rectangles represent complete rater agreement, and the gray rectangles represent partial agreement, i.e., adjacent ratings, such as Never vs. Rarely, or Rarely vs. Sometimes.

The weighted Bangdiwala statistic for rater 2 agreement for large game was B = 0.85, and for rater 3 was B = 0.81. See Fig S2.

Fig. S2: Large game agreement plots for the original ratings (x-axis) vs. rater 2 (y-axis, left) and rater 3 (y-axis, right). The interpretation is as in Fig S1.

Comparing the independent re-coding of rater 3 with the original re-coding, we identified 10 societies for which there were substantial discrepancies, i.e., a difference of two or more on our ordinal scale (e.g., a coding of “Never” vs. “Sometimes”). These major discrepancies were resolved and the original recode data set updated, which is what we report.

## S3 Figure

Fig. S3: Plot of the Anderson et al. (2023) data (left), and our recoding (right). The selection of societies in Anderson et al. (2023) is heavily biased, so neither plot is representative of female hunting in foraging societies.

## S1 Table

[D-PLACE comparison](https://docs.google.com/spreadsheets/d/12-gORpmwOLrptdyIdtqr6JkoDtzsM6p6bPpPW-RJQGA/edit#gid=88624079)

## S2 Table

Recoding Anderson

## S1 Data and code

**Re-coded data and analysis scripts** https://github.com/grasshoppermouse/anderson_critique

## Acknowledgments

We thank Polly Wiessner and Elizabeth Cashdan for their helpful comments.

